# Computational rewiring of allosteric pathways reprograms GPCR selective responses to ligands

**DOI:** 10.1101/2022.03.30.486353

**Authors:** D. Keri, M. Hijazi, A. Oggier, P. Barth

## Abstract

G-protein-coupled receptors (GPCRs) are the largest class of cell surface receptors and drug targets, and respond to a wide variety of chemical stimuli to activate diverse cellular functions. Understanding and predicting how ligand binding triggers a specific signaling response is critical for drug discovery and design but remains a major challenge. Here, computational design of GPCR allosteric functions is used to uncover the mechanistic relationships between agonist ligand chemistry, receptor sequence, structure, dynamics and allosteric signaling in the dopamine D2 receptor. Designed gain of function D2 variants for dopamine displayed very divergent G-protein signaling responses to other ligand agonists that strongly correlated with ligand structural similarity. Consistent with these observations, computational analysis revealed distinct topologies of allosteric signal transduction pathways for each ligand-bound D2 pair that were perturbed differently by the designs. We leveraged these findings by rewiring ligand-specific pathways and designed receptors with highly selective ligand responses. Overall, our study suggests that distinct ligand agonists can activate a given signaling effector through specific “allosteric activator” moieties that engage partially independent signal transmission networks in GPCRs. The results provide a mechanistic framework for understanding and predicting the impact of sequence polymorphism on receptor pharmacology, informing selective drug design and rationally designing receptors with highly selective ligand responses for basic and therapeutic applications.

## Introduction

G protein-coupled receptors (GPCRs) are 7 transmembrane signaling proteins that enable cells ability to communicate with the extracellular (EC) environment. With over 800 human GPCRs, these receptors comprise roughly 3% of the human genome. They sense a wide range of signals including elementary particles, small molecules, peptides and lipids, and translate these extracellular stimulations into diverse physiologically important intracellular responses^1–4^. As such, they represent an important class of drug targets and close to 30% of all FDA approved drugs target GPCRs^5^. In addition to the large number of functionally diverse GPCRs expressed in most body’s tissues, each receptor often functions as a signaling hub, able to sense distinct ligands and elicit different ligand-specific intracellular responses. Over the past decade, structural and molecular studies have begun to illuminate the structural underpinnings of ligand binding and stabilization of GPCRs^6–9^. However, our molecular mechanistic understanding of how binding of distinct ligands leads to differential receptor signaling (i.e. functional selectivity or biased agonism) remains limited^10–13^, owing to the complexity of signaling transduction processes across lipid membrane.

GPCRs translate extracellular stimuli into intracellular functions through long-range allosteric signal transduction across lipid membranes. Allostery enables ligand-induced changes in protein structure and dynamics to be efficiently transmitted to distant sites and is a widespread regulation mechanism of protein function^14–16^. Owing to the lack of high-resolution dynamical measurements on GPCRs, how allostery is encoded into receptor sequence, structure and dynamics is not well understood. Computational studies using sequence co-evolution inference or molecular dynamics simulation can detect networks of functionally and dynamically coupled receptor residues that may provide efficient communication pathways^17–20^. However, how distinct ligands engage these networks to elicit precise and selective signaling responses remains elusive.

In this study, we aimed to better understand the relationships between ligand agonist structure and chemistry on one side, and receptor sequence, structure, dynamics and allosteric signaling on the other side using the dopamine D2 receptor as a model system. Several ligand-bound D2 structures have been solved recently and D2 functions are regulated by many well characterized drug ligand molecules^21–24^. Additionally, we recently developed a computational approach for designing GPCRs with altered signaling responses through engineered allosteric microswitches^25^. Using the method, we designed novel D2 receptors with enhanced sensitivity and G-protein responses for the strong dopamine and weak serotonin agonist ligands that provide one of the foundations for this study.

## Results

In a previous study, we had identified a class of allosteric sites (“allosteric propagators”) that connect highly conserved microswitches into fully wired allosteric pathways. We were able to fine-tune GPCR signaling responses through novel allosteric “propagator” microswitches designed on several TMHs, which suggested the existence of multiple allosteric signal transduction pathways running through the TM region of the receptor^25^. The designed microswitches at propagator sites enhanced the sensitivity to both dopamine and serotonin, implying that receptor responses to both ligands may involve the same path. Since signal transduction pathways connect the extracellular ligand to the intracellular effector binding sites, we reasoned that ligands with similar structure should engage similar paths through overlapping contacts with the receptor. Conversely, structurally distinct agonists that bind to the receptor through different “allosteric activator” chemical groups should involve alternative pathways and therefore be sensitive to different allosteric “propagators”.

To test this hypothesis, we sought to explore the mechanistic relationships between agonist ligand chemistry and receptor signaling through the atomic-resolution mapping of allosteric pathways and quantification of signal transductions in ligand-receptor systems (**Fig. 1**). We took a multi-disciplinary approach combining ligand structure clustering for mapping chemical space, molecular dynamics simulation (MD) for allosteric pathway discovery, computational protein design for reprogramming allosteric responses and a battery of cell signaling assays for measurement of receptor responses to multiple ligands (**Fig. 1a**). If we understand how ligands activate a receptor through engagement of specific allosteric pathways, we should be able to rationally rewire these paths and design novel selective ligand-GPCR responses (**Fig. 1b**).

**Figure 1.**
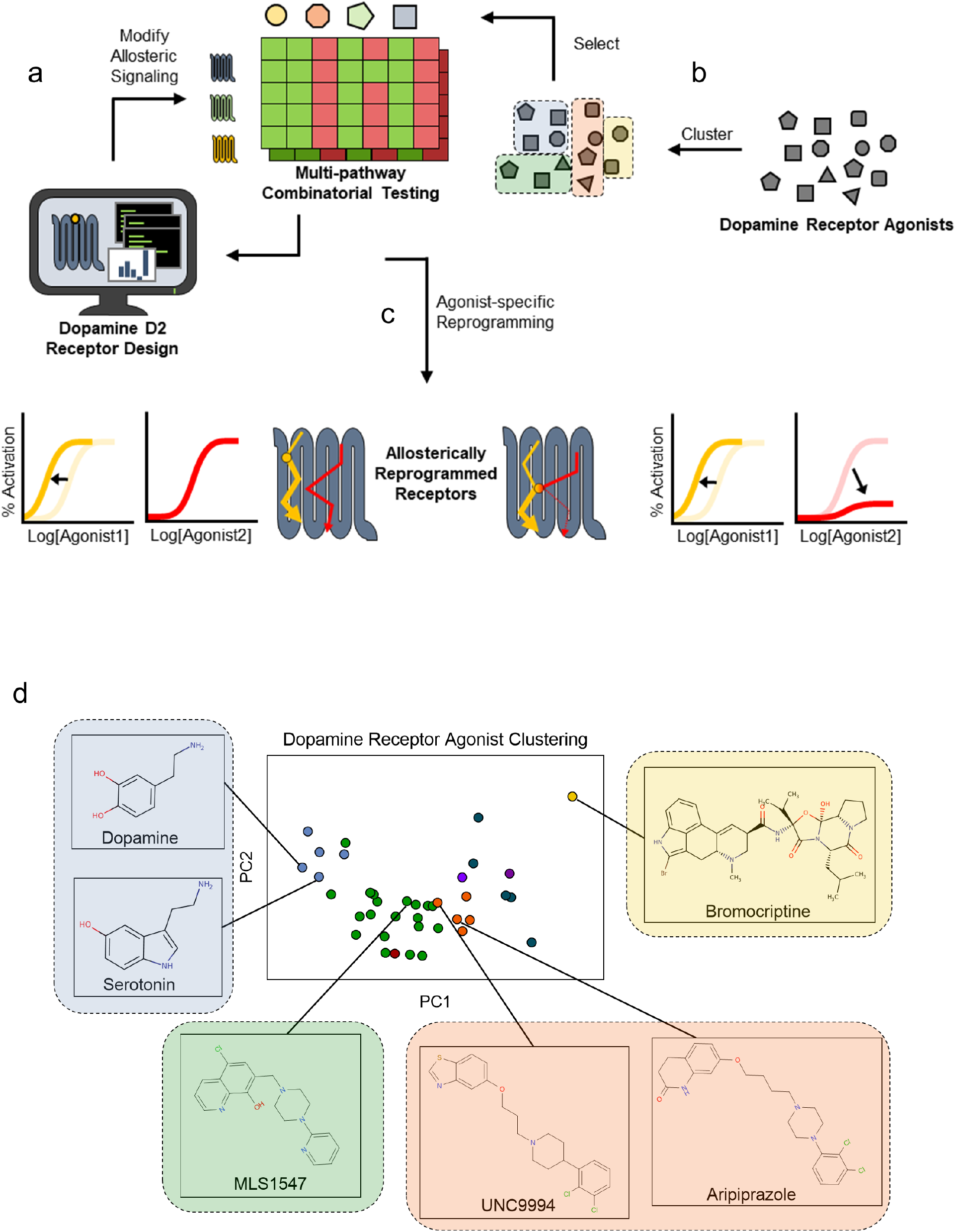
Outline of computational and experimental workflow. a. Computationally designed receptors with predicted changes in allosteric signaling are tested against a set of structurally diverse agonists using G protein activation assays, in an iterative process. The first round of designs is agonist-agnostic while subsequent rounds utilize ligand-docked receptor models. b. Structural clustering and representative ligand selection. c. Design selection based on ligand-specific functional output. d. Principal component analysis plot of clustered dopamine receptor agonists. Highlighted agonists were chosen for experimental testing and are colored based on cluster.

We selected the dopamine D2 receptor as a system of study because it performs critical neurological functions^26,27^, has been structurally characterized in several signaling states and is regulated by numerous partial and full ligand agonists. We first analyzed the structural similarity of a comprehensive set of 39 characterized D2 agonists using standard structural clustering approaches and identified 4 major clusters (**Fig. 1c, methods**). Consistent with our expectations, serotonin (SE) and dopamine (DA) adopt very similar chemical structure and therefore belong to the same cluster. D2 agonists from other clusters are often much larger than DA and SE, populate distinct chemical subspaces and can presumably contact the receptor through multiple additional sites.

To investigate the relationship between ligand chemical space and long-range allosteric signaling responses, we first assessed how several allosteric propagator microswitches previously-designed in the TM core of D2 modulate the receptor response to structurally-distinct ligands (**Fig. 2a**). Because these sites are far away from the ligand binding site, they can probe long-range interactions connecting ligands and allosteric pathways running though the receptor structure. We selected representative members of the 4 ligand clusters and measured in vitro the D2-mediated activation of the G protein Gi2 upon ligand stimulus using HEK reporter cell lines stably expressing a Trp channel (**Fig. 2b**). Four designed D2 receptors incorporating distinct allosteric propagator microswitches at TMHs 5 and 6 (**Fig. 2c**) were transiently expressed in stable HEK cells and incubated with the following 6 agonist ligands: the full agonists dopamine (DA), bromocriptine (BRC), and the partial agonists serotonin (SE), aripiprazole (AR), MLS1547 (MLS) and UNC9994 (UNC). Dose titrations revealed very distinct effects of the designed microswitches on the assayed ligands. The large (i.e. from 1 to 2 orders of magnitude) increases in dopamine and serotonin potency and efficacy (**Fig. 2d,e**) were not observed for the other ligands (**Fig. 2f-i**). While the designed microswitches still behaved as gain of function for the partial agonists, smaller increases in efficacy and potency were observed. By contrast, opposite effects were measured for the full agonist bromocriptine that lost efficacy and potency for all but the T5.54M (residues are numbered according to the Ballesteros-Weinstein numbering scheme^28^) microswitch which behaved similarly to WT (**Fig. 2f**).

**Figure 2.**
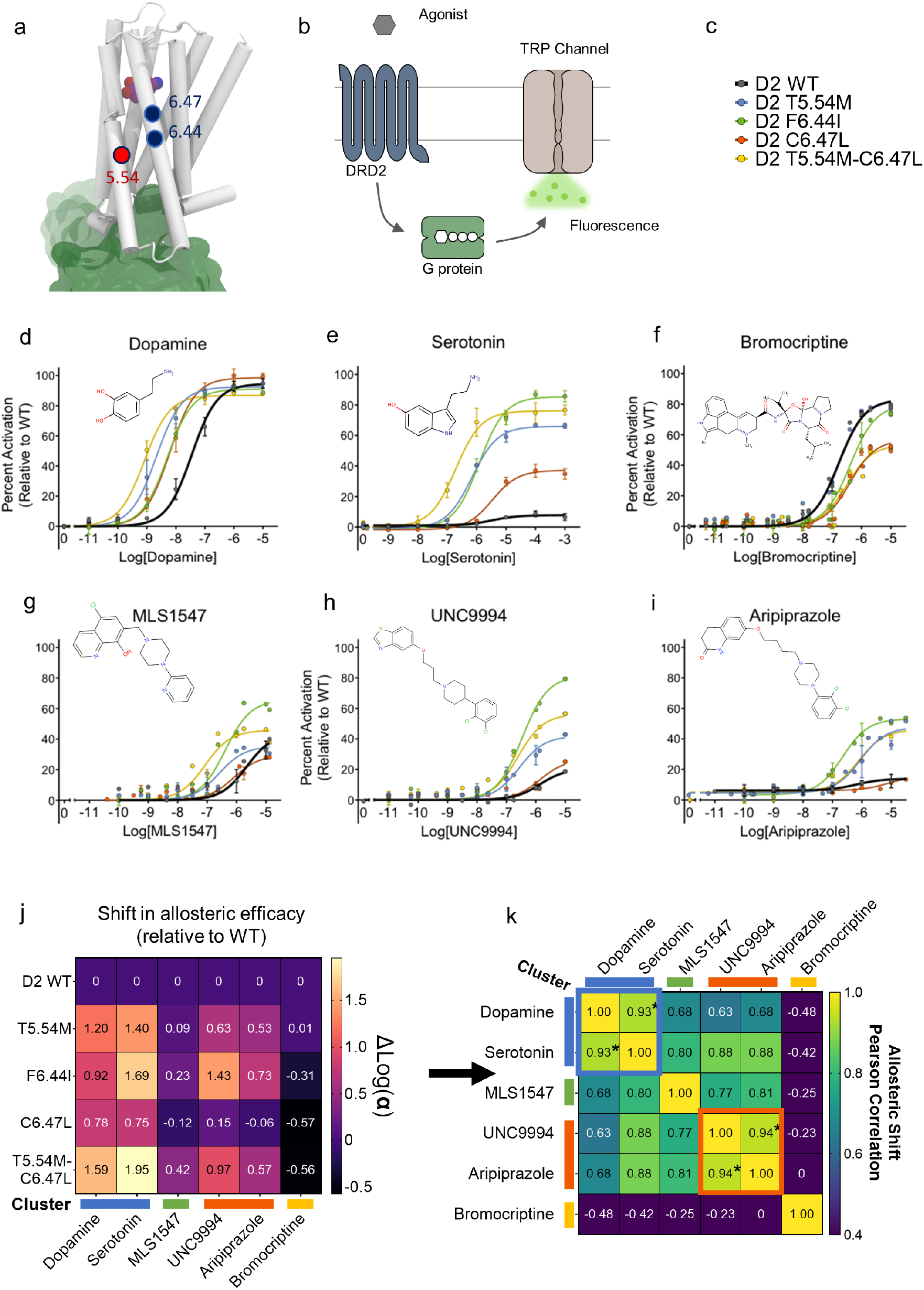
Initial workflow input and first round functional output. a. Positions of the initial ligand-agnostic designs on the dopamine D2 receptor structure. b. Diagram of the fluorescence-based Transient receptor potential (TRP) channel assay for the detection of G protein Gi activation. c. Designed D2 variants. d-i. Agonist-dependent G protein activation normalized to D2 WT dopamine response. Values are as mean ± standard error of the mean (SEM). n = 3 j. Combinatorial shifts in allosteric efficacy based on the allosteric two-state model (ATSM). k. Pearson correlations of allosteric efficacy shifts between agonists for all tested dopamine D2 receptor designs. *p < 0.05.

To better understand the origin of these surprising observations, we interpreted the results within the framework of a two state allosteric model and quantified the change in allosteric coupling α underlying the shifts in ligand efficacy (**Fig. 2j**). Here the parameter α describes the efficiency with which a ligand engages and stabilizes the receptor active state. Remarkably, each design was characterized by a specific allosteric coupling signature across the spectrum of tested ligands. T5.54M displayed strong, mild and no gain of function effects for DA/SE, MLS/UNC/AR, and BRC, respectively. On the other hand, F6.44I displayed strong, mild gain of function and loss of function effects for DA/SE/UNC/AR, MLS, and BRC, respectively. The largest differences were observed for the double switch T5.54M-C6.47L with very strong, strong, very mild gain of function and loss of function effects for DA/SE, UNC/AR, MLS, and BRC, respectively. Interestingly, specific designed effects tend to be similar for ligands from the same cluster and could be classified into 4 distinct classes; i.e. DA/SE, UNC/AR, MLS, BRC (**Fig. 2k**). The results suggest that, while each allosteric propagator microswitch has the potential to modulate ligand efficacy differently, the effects correlate with how ligand activate the receptor and trigger allosteric pathways. Importantly, the allosteric coupling profiles could not be explained by the differences in efficacy of the ligands for the WT receptor as DA and BRC are both full agonists and SE is a very weak agonist for D2.

To better understand the origin of these observations and further dissect the allosteric determinants underpinning the signaling responses to DA and BRC, we sought to design additional allosteric propagator microswitches in other regions of the receptor. We carried out calculations using our allosteric design pipeline^25^ and scanned TM regions involving TMHs 2, 3 and 4 that do not undergo significant conformational changes upon receptor activation and were not included in our previous study. The simulations identified several candidate microswitches that were predicted to selectively increase the allosteric coupling in the active state and enhance the response to ligand agonists (**Fig. 3a-c**). Among these, the microswitch L3.41H considerably improved the receptor sensitivity to DA, boosting the ligand potency by close to 15 fold (**Fig. 3b**). Unlike the other designed microswitches, L3.41H was also a gain of function variant for BRC (**Fig. 3c**). Overall, while the designed microswitches tested so far all enhanced DA potency, we observed much more specific effects for BRC that tend to depend on the location of the propagator. Indeed, microswitches on TM6, TM5 and TM3 were loss of function, neutral or gain of function for BRC, respectively.

**Figure 3.**
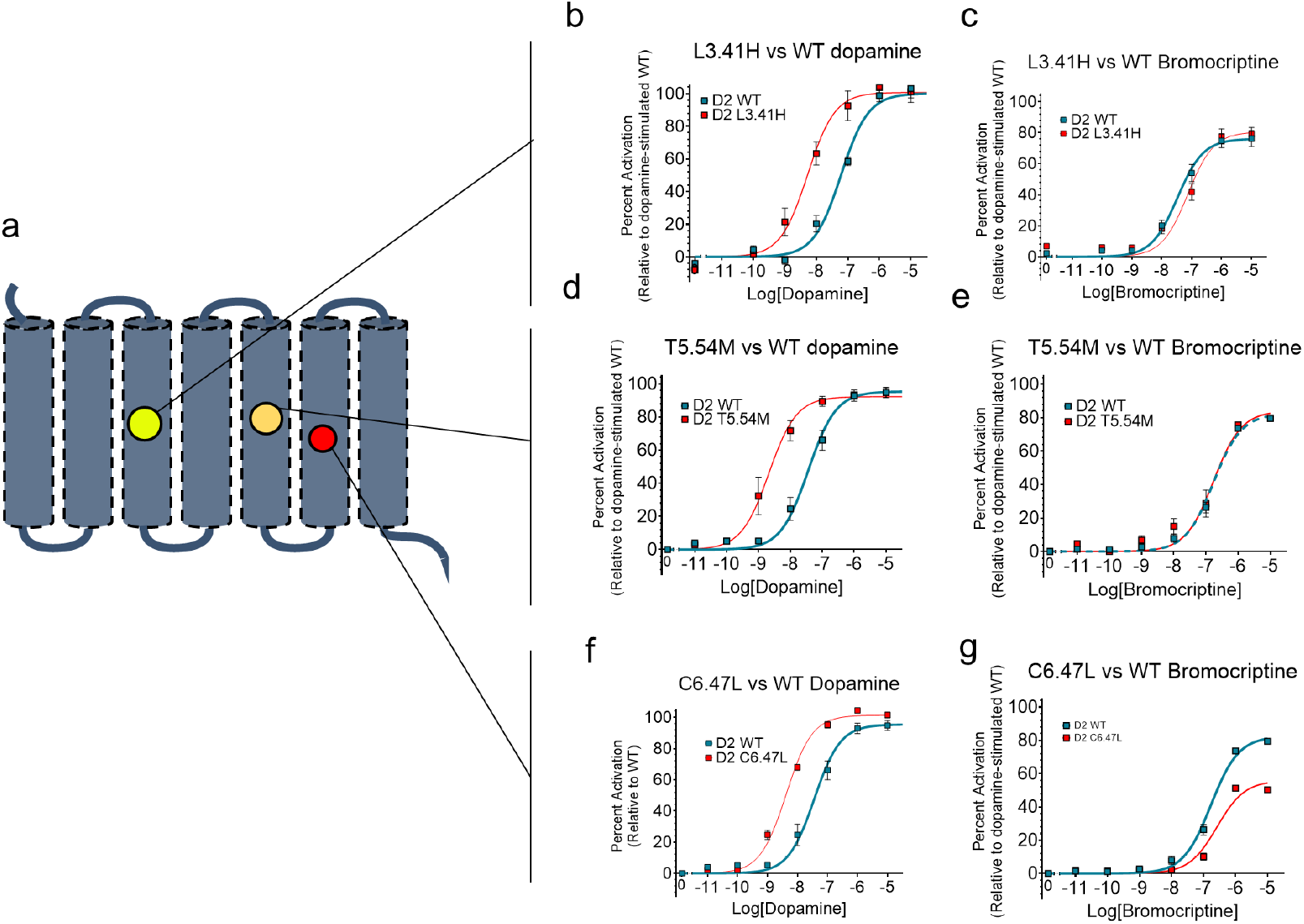
Highlight of divergent agonist response shifts. a. Diagram of the dopamine D2 receptor showing design positions resulting in higher dopamine sensitivity with diverse bromocriptine response shifts. b-g. TRP channel assay dose response curves with dopamine (left column) and bromocriptine (right column). Points shown are mean ± SEM. n ≥ 3.

These observations suggested the existence of several allosteric pathways running through distinct TMH interfaces that would be potentiated differently by the two full agonists to activate Gi, hence prompting us to investigate the DA and BRC-bound receptor structures. We first analyzed the ligand-receptor short-range contacts. Consistent with the large differences in ligand size, BRC displayed 100% more contacts with the orthosteric site than DA (**Fig. 4a-d**). While most D2 residues interacting with DA were found to also contact BRC directly, 12 D2 residues on TMH 3, ECL2, TMHs 6 and 7 were contacting BRC only (**Fig. 4e**). To investigate the strength and persistence of these contacts, we relaxed each ligand bound D2 structure in explicit lipid bilayer using Molecular Dynamics over 1.25 μs simulation time. Most contacts inferred from either the cryoEM BRC-D2 structure or the DA-D2 model were persistent in our simulations. Consistent with a higher density of receptor-ligand contacts, the BRC-D2 structures displayed significantly lower Root mean square fluctuations (RMSF) than DA-D2 structures especially in regions directly in contact with the ligand.

**Figure 4.**
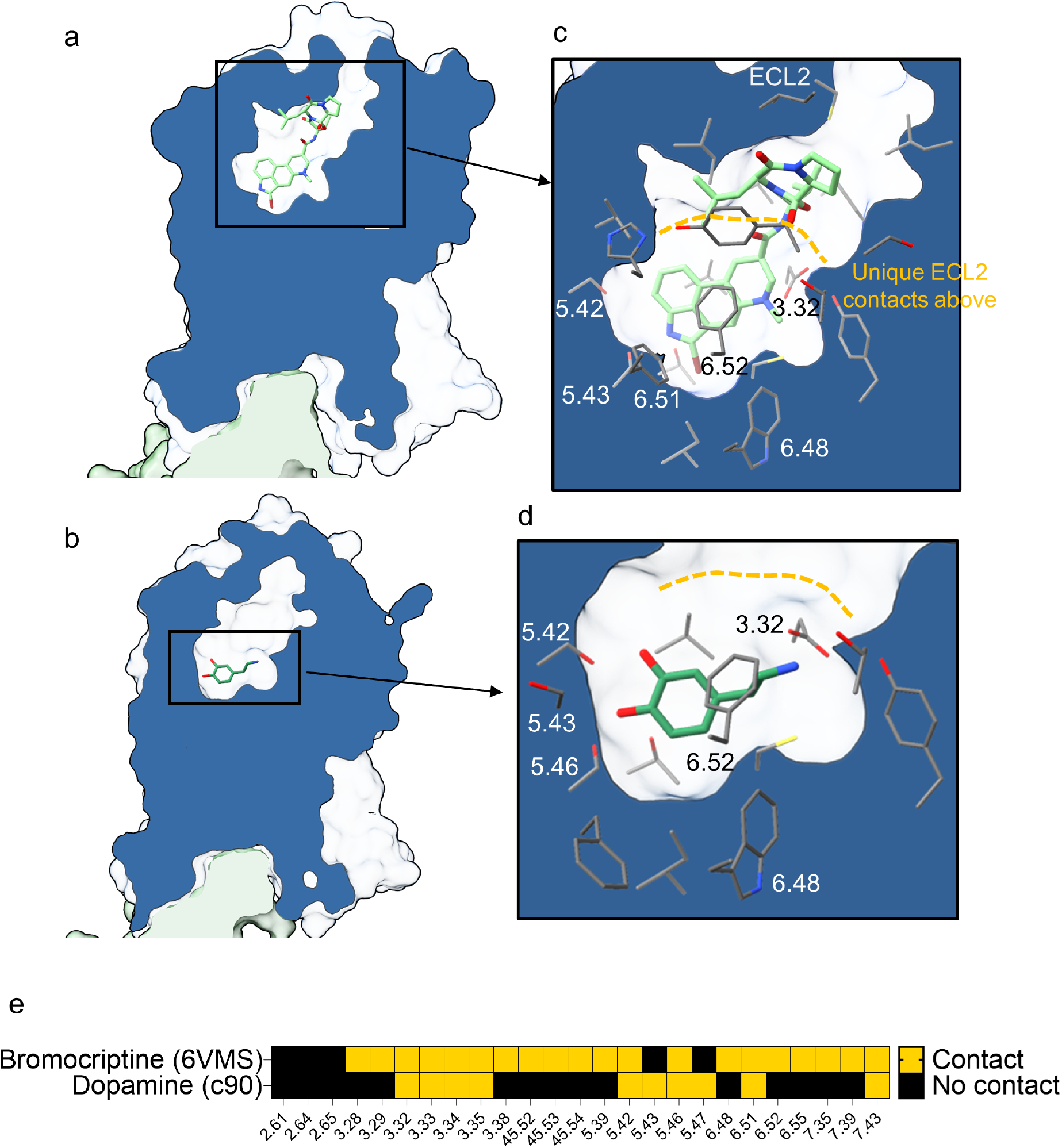
Ligand-receptor short-range contacts. Bromocriptine (a, c.) and dopamine (b, d.) ligand binding pockets shown in context of the full D2 receptor (a, b.) and close-up of the ligand binding pose and contacting residues (c, d.). Bromocriptine binding pose was extracted from Gi-bound active state D2 structure (PDB code 6VMS) while dopamine’s binding pose was extracted from a consensus ligand docking model (see methods). e. Contact map showing residues contacting bromocriptine and dopamine respectively. Residues are numbered according to the Ballesteros-Weinstein numbering scheme.

To assess how the differences in rigidity affect the allosteric signaling pathways running through the receptors, we analyzed the dynamic fluctuations extracted from the all atom MD trajectories and calculated the Mutual Information (MI) between all pairs of residues. MI characterizes the amount of information exchanged between two residues and provides a measure of mechanical coupling between 2 sites. Allosteric signals are predicted to be best propagated by residues that exchange the most information^29^. Hence, we identified the subset of residues with the highest MI that create contiguous networks connecting the orthosteric to the intracellular binding sites (**Methods**)^17,30^. We clustered pathways running through the same residues into pipelines and ranked them according to path densities.

From the top 10 ranked pipelines, we analyzed those connecting the extracellular to the intracellular regions for the following ligand-receptor pairs: DA-D2_WT_; DA-D2_T5.54M-C6.47L_; DA-D2_L3.41G_; DA-D2_L6.41M_; DA-D2_F6.44M_; BRC-D2_WT_; BRC-D2_T5.54M-C6.47L_; BRC-D2_L3.41G_; BRC-D2_L6.41M_; BRC-D2_F6.44M_ (**Fig. 5a,b**). Owing to divergent DA and BRC binding contacts, the major paths in D2_WT_ were mainly initiated by different residues located on the extracellular tips of TMHs 5, 6, 7 for DA and TMHs 5, 6 and ECL2 for BRC. Except for a major common path running through TMH 5, the topology and predominance of the paths were also quite distinct between the 2 ligand-bound D2_WT_ structures. For example, top-ranked paths running all the way through TMH 6 were only observed in the BRC-D2WT structure. While BRC and DA bound structures display a similar path running through TMH 7 to H8, it was more prominent in the DA bound receptor.

**Figure 5.**
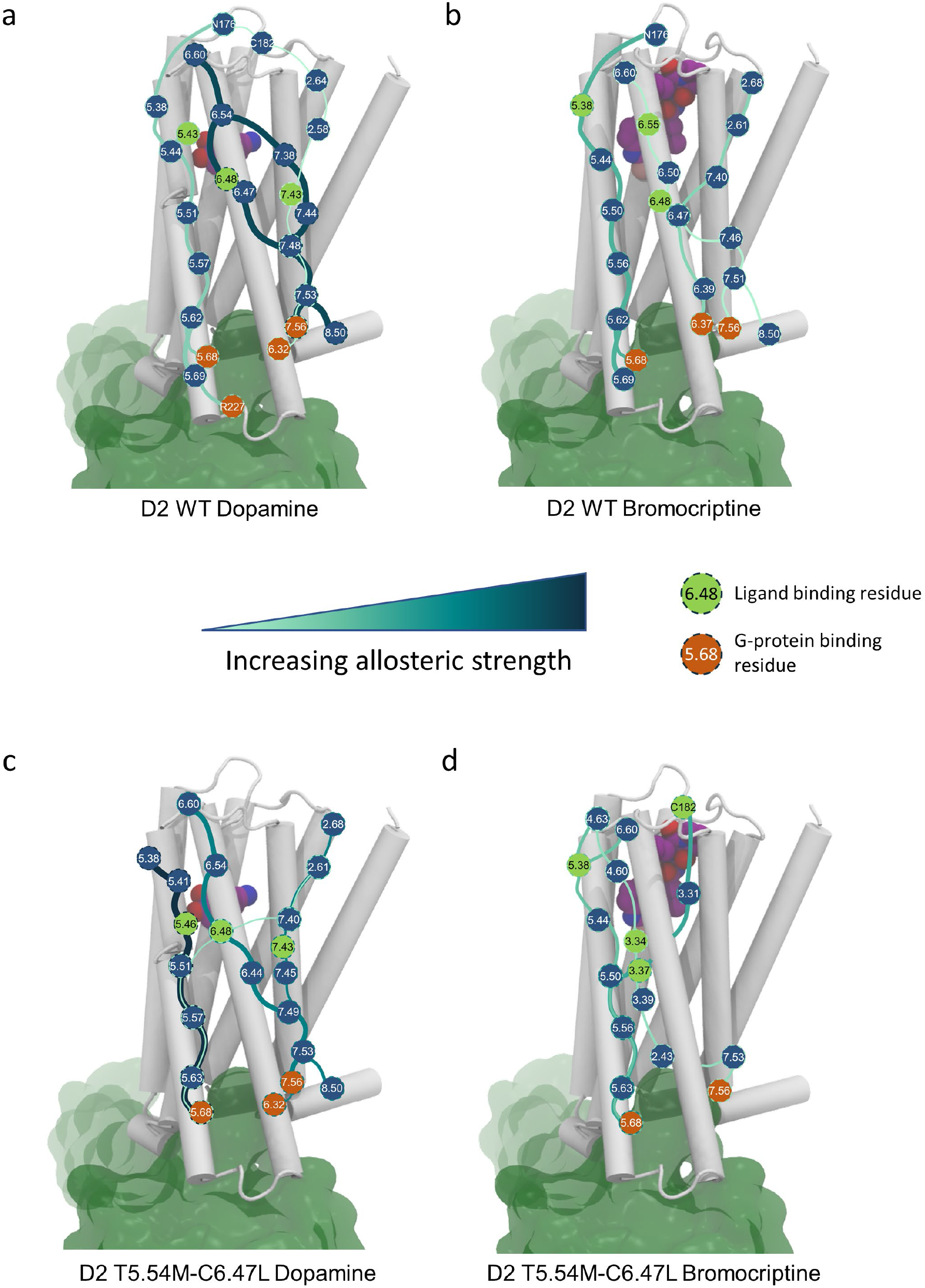
Allosteric pipelines extracted from all-atom molecular dynamics simulations for a. WT D2 bound to dopamine b. WT D2 bound to bromocriptine c. T5.54M-C6.47L D2 bound to dopamine and d. T5.54M-C6.47L D2 bound to bromocriptine. Bound ligand is shown in purple space filling representation and bound G-protein is shown in green surface representation. Allosteric pipelines visualized represent the top scoring pipelines that connect the ligand binding region to the G-protein binding region, with the color shade and thickness of pipelines representing allosteric communication strength, which is measured as the path density (see methods). Residues are numbered according to the Ballesteros–Weinstein numbering scheme.

Unexpectedly, the designed allosteric propagator switches considerably rewired the pathways initiated by both ligands (**Fig. 5**). For example, the path running through TMH 5 in DA-D2_WT_ was greatly enhanced in DA-D2_MUT_, with a new allosteric pipeline connecting to TMH 5 through TMHs 6 and 7. We also observed new path connectivities in the neighborhood of the designed switches, particularly at the interface between TMHs 6 and 7 in DA-D2_MUT_. However, the largest changes were observed for the receptor bound to BRC (**Fig. 5c,d**). Two prominent paths running through TMHs 6 and 7 in BRC-D2_WT_ were substantially weakened in BRC-D2_MUT_ and disappeared from the top ranking pipelines. Instead, a new path emerged in the designed structure initiated by residues on TMHs 3 and 4, running through TMH 3 and branching out to reach TMH 7 and H8 on the intracellular side. A large body of structural evidence indicate that TMHs 6 and 7 movements are critical for opening the receptor intracellular side to G-protein binding. Hence, paths that mechanically couple agonist ligands to the intracellular tips of TMHs 6 and 7 are expected to be key determinants of receptor’s ability to recruit and activate G proteins. Our calculations suggest that the designed microswitches significantly weakened the coupling of BRC to G protein binding and should therefore decrease the ligand efficacy, which is consistent with our experimental observations.

If DA and BRC exploit distinct pathways to activate the same signaling effector, we should be able to create D2 receptors with high ligand selectivity by reprogramming long-range communication along ligand-specific allosteric channels. We first identified the main allosteric sites through which DA or BRC-specific paths were predicted to run through and defined DA or BRC-selective allosteric hubs. We then applied our computational allosteric design approach to select *in silico* novel combinations of amino-acids at these hubs predicted to enhance or weaken the allosteric coupling mediated by the ligand selective paths. Overall, since our first generation of designs were primarily selected to elicit stronger DA responses regardless of the effects on BRC, our goal was to engineer either gain of function variants for BRC or receptors with more ligand selective responses. We targeted a wide variety of sites in the TM core of the receptor, including TMHs 3 and 4 that were not covered in our previous study (**Fig.6a**). We validated our selected designs by measuring ligand-stimulated Gi-activation responses in Trp-HEK cells as described above. Consistent with our intentions, we were able to engineer D2 receptors with a much higher variation in response to BRC when compared to our first generation of designs (**Fig.6b-e**). In particular, designed microswitches at sites 3.41 and 6.41 enhanced BRC efficacy by up to 36% and 48%, respectively (**Fig.6c, d**). Remarkably, these effects were highly selective for BRC in the 3.41G design which did not show any difference in DA responses as compared to WT (**Fig.6c**). Conversely, sequence changes at the 4.46 hub had little effect on DA but considerably decreased the response to BRC. For example, no Gi activation signals by the I4.46N design could be detected even at submillimolar concentrations of BRC. Lastly, F6.44M strongly enhanced the sensitivity for DA while suppressing all Gi-mediated response to BRC. This receptor achieved a very high level of ligand selectivity and, together with the other designs, demonstrates that the rational rewiring of selective allosteric pathways enables the fine-tuning and control of ligand mediated receptor functions.

**Figure 6.**
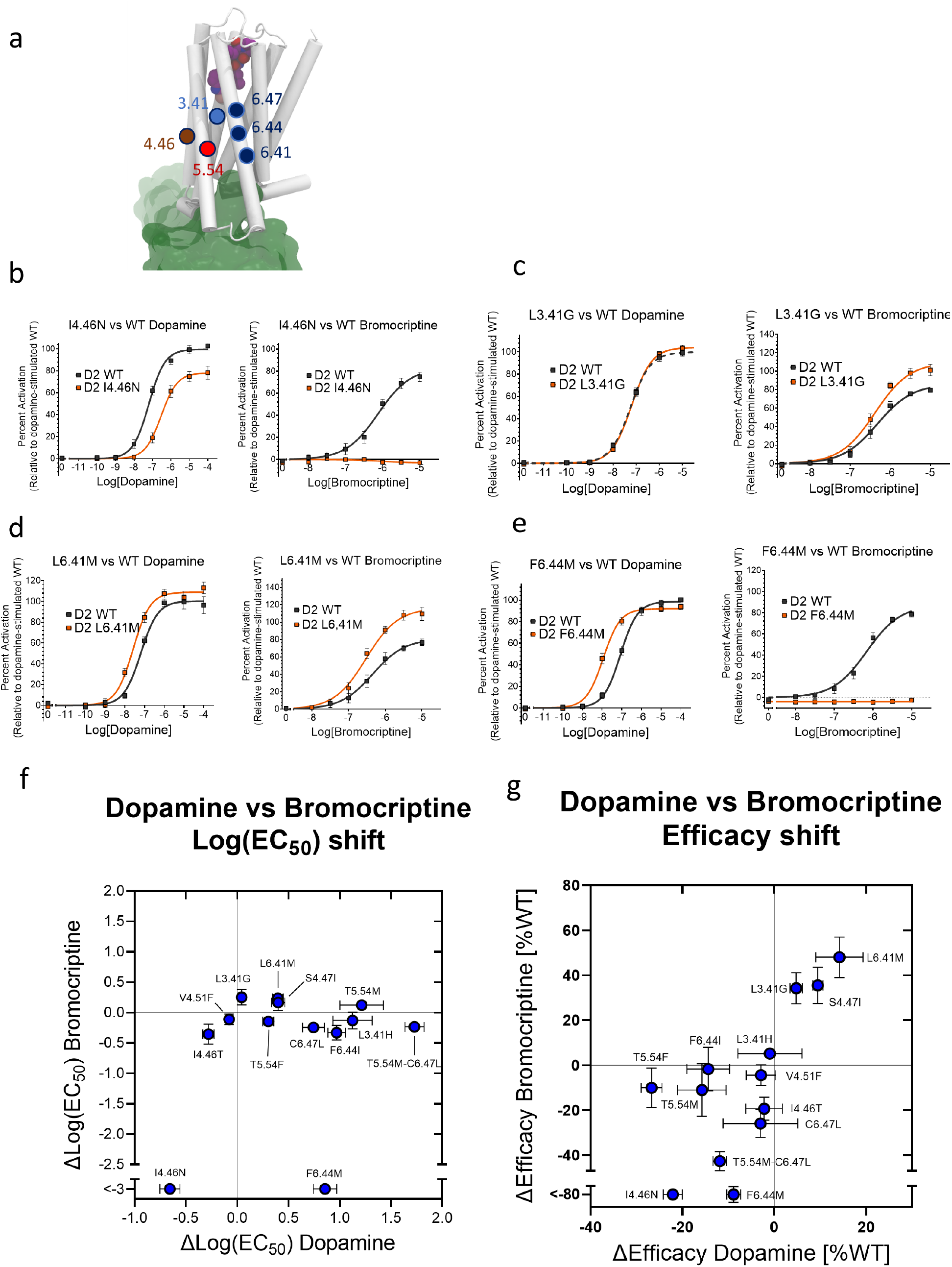
Agonist-specific allosteric pathway reprogramming. A new paradigm for GPCR agonist-specific signaling. Different agonists utilize partially distinct allosteric pathways to activate the receptor. These allosteric pathways can be independently manipulated via mutation. a. Positions of the ligand-specific designs on the dopamine D2 receptor structure. b-e. Dopamine (left column) and bromocriptine (right column) dose response curves for I4.46N, L3.41G, L6.41M, and F6.44M. Values are mean ± SEM. n ≥ 3. f,g. Scatter plots of designs with altered dopamine or bromocriptine signaling plotted according to agonist-specific shifts in sensitivity (Log(EC50), f) and efficacy (maximal response, g).

## Conclusion

In this study, we explored the relationships between agonist ligand chemical space and D2 signaling responses through the lens of receptor conformational dynamics, allostery and protein design. We found that allosteric microswitches designed in the receptor TM core to enhance ligand-mediated Gi activation had gain of function effects that were very specific to ligand agonists. A strong correlation was observed between ligand structural similarity and the functional shifts measured for ligand-D2 pairs and prompted us to further investigate the two full agonist ligands (DA and BRC) that displayed the most divergent Gi-mediated responses to designed microswitches. An in-depth analysis by MD simulations revealed a distinct topology of allosteric pathways connecting each ligand to the intracellular Gi binding site and different path perturbation and rewiring upon designed microswitches that were consistent with the measured changes in ligand responses. To validate these findings, we designed a second generation of allosterically reprogrammed receptors where predicted ligand-selective allosteric paths were strengthened or weakened. As intended by the calculations, the designs displayed strong ligand-selective Gi activation shifts, including a highly sensitive receptor for DA that completely lost its ability to respond to BRC. Overall, our study suggests that distinct ligand agonists can activate a given signaling pathway through specific “allosteric activator” moieties that engage partially independent allosteric pathways running through the receptor. Overall, our ability to rationally design receptor sequence variants with fine-tuned signaling responses to specific ligands paves the way for engineering very selective ligand biosensors and predicting the effects of receptor polymorphisms on drug pharmacology for enhanced personalized medicine.

## Methods

### D2 ligand clustering

A Dopamine D2 receptor ligand list was obtained from the CHEMBL database (https://www.ebi.ac.uk/chembl/) and only agonists were kept. The physicochemical properties of all agonists were summarized using the JOELib cheminformatics library. The resulting data were clustered using density-based clustering (DBSCAN)^31^ and finally dimensionality were reduced using principal component analysis (PCA).

### TRP channel cell-based assay

HEK-293 cells stably expressing the TrpC4β channel (generously provided by Dr. Michael X. Zhu) were maintained in DMEM supplemented with 10% fetal bovine serum (FBS) and 50μg/mL geneticin as a selective antibiotic and grown at 37°C and 5% CO_2_. FMP2 FLIPR Membrane potential assays (Molecular Devices) were performed as previously described^25^. Briefly, the assay relies on the detection of a membrane-permeable fluorescent dye coupled to a non-permeable quencher. The non-selective cation TRP channel changes the membrane potential upon activation by Gi, which enables the selective influx of the dye. 24 hours prior to the assay, 150,000 cells/well are seeded into clear-bottom 96-well plates and are reverse transfected with an optimized quantity of receptor DNA (present in the pcDNA3.1+ vector) and 0.5μL Lipofectamine 2000 per well. Prior to reading the transfected plates, the media is removed and the FLIPR dye is applied. The relevant drug is transferred into the plates during plate reading on a Flexstation3 multi-mode plate reader and changes in fluorescence (emission at 535nm, excitation at 565nm) are measured for a period of a maximum of 4 minutes.

### Enzyme-linked Immunosorbent Assay (ELISA)

To ensure all receptor variants express at a level similar to the dopamine D2 WT receptor control, ELISA’s are performed against the 3xHA N-terminal tag present on each receptor variant. For each of the aforementioned cell-based assays, an accompanying ELISA plate is also reverse transfected in parallel using the same conditions as the main assay plate. On the day of the assay, the media is removed from the wells and the cells are fixated via a 4% paraformaldehyde (PFA, Electron Microscopy Sciences) solution incubation for 10 minutes. Fixation is followed by a 2% bovine serum albumin (BSA, Sigma Aldrich) solution incubation, anti-HA mouse primary antibody (Santa Cruz Biotech) incubation, and an anti-mouse secondary HRP-linked antibody incubation (Thermo Fisher); each for a period of 1 hour with three PBS washes between each step. SuperSignal chemiluminescent substrate (Thermo Fisher) is added to each well and plates are incubated for 5 minutes before a luminescent reading on a Flexstation3 plate reader. Mock-transfected wells are used to determine and subtract the baseline signal from the remaining wells.

### Ligand docking

Dopamine docking onto the solved dopamine D2 receptor active state structure (6VMS^23^) was accomplished by the established IPhold Rosetta protocol^32^. Only the receptor chain was used during all steps; heteroatoms and additional chains were removed. The overall protocol consists of sequential coarse-grained docking coupled with structural relax, decoy clustering, high resolution docking, and ligand clustering steps. During the first docking and relax step 10,000 decoys are generated, of which the lowest 10% scoring are used in the subsequent structure clustering step to diversify target receptor conformation. High resolution docking was performed on the cluster centers of the largest 6 clusters, again generating 10,000 decoys for each cluster center model and only using the lowest 10% scoring decoys for subsequent analysis. An additional filter was added to only keep the lowest 50% scoring decoys in terms of interface energy (interface_delta). The remaining decoys were used for ligand binding mode clustering using a DBSCAN algorithm with ligand heavy atom coordinates as the input. The largest binding mode cluster was designated as the putative native binding mode and used for further analyses.

### *In silico* mutagenesis

Dopamine D2 variant models were generated using RosettaMembrane^33^, a Rosetta-based protein structure prediction software utilizing a Monte Carlo gradient descent energy minimization algorithm enhanced with an anisotropic implicit membrane scoring functionality. The recently released active-state dopamine D2 receptor structure (6VMS^23^) served as the initial starting structure. All heteroatoms and non-receptor or G-protein chains were removed from the starting structure. Residues of interest were mutated and adjacent residues within 5Å were subjected to alternating cycles of sidechain repacking and backbone relaxation through Rosetta’s Monte Carlo-based energy minimization algorithm. 200 decoys were generated per design to ensure score convergence. The lowest scoring decoys were used for all subsequent analyses.

### Molecular dynamics simulations and allosteric pathway predictions

Obtaining allosteric pipelines that are used in design requires two prerequisite steps: the first is running molecular dynamics simulations and the second is calculation of mutual information from the simulations.

The starting structures used for MD simulations are: 1-BRC bound 6VMS with the last 20 residues of the C-terminal helix of Gi and the sequence re-mutated back to WT, and 2-the DA bound docked model based on 6VMS, in addition to the mutated (T5.54M-C6.47L) 3-DA-bound and 4-BRC-bound generated based on the WT structures using Rosetta^33^. The receptor-ligand-helix complex was inserted in a 90 × 90 Å^2^ POPC lipid bilayer solvated by 22.5 Å layer of water above and below the bilayer with 0.15 M of *Na*^+^ and *Cl*^−^ ions using CHARMM-GUI bilayer builder^34,35^. Parameters for the two ligands (dopamine and bromocriptine) were generated using CGenFF^36^. Simulations were performed with GROMACS 2019.4 with CHARMM36 forcefield^37^ in an *NPT* ensemble at 310K and 1 bar using a velocity rescaling thermostat (with a relaxation time of 0.1 *ps*) and Parrinello-Rahman barostat (with semi-isotropic coupling at a relaxation time of 5 *ps*) respectively. Equations of motion were integrated with a timestep of 2 *fs* using a leap-frog algorithm.

Mutual information was then calculated from torsional angles extracted every 100 ps from the simulations by histogramming the data and then calculating probabilities followed by 1^st^ order and 2^nd^ order entropies and mutual information, as described by Killian et al.^38^.

After that, mutual information was fed into Allosteer package to calculate allosteric communication pipelines^17,30^. In short, Allosteer calculates the shortest pathways starting from extra-cellular (EC) region passing through the ligand binding region and reaching all the way to G-protein binding region that maximize mutual information. Overlapping allosteric pathways are clustered into allosteric communication pipelines, and the strength of a pipeline is the number of pathways passing through it. We considered the top 10 ranking pipelines for our analysis, and then chose the pipelines from the top 10 ranking that contact both the ligand and the G-protein (Fig. 5). Allosteric hubs are defined as the residues with the largest number of allosteric pathways passing through them. Ligand binding residues are defined as residues in contact with the ligand (DA or BRC) for at least 55% of total simulation time, while G-protein binding residues were extracted from bromocriptine bound D2 structure PDB code 6VMS. Ligand or G-protein and receptor residue are considered in contact if their heavy atoms are within 5Å.

### Allosteric pathway coupling strength calculation

Global allosteric coupling through receptor structures was estimated as previously published^25^ using normal mode analysis. To calculate a “coupling” value for each receptor variant, the list of allosteric hubs considered varies. Residues corresponding to an established list^17^ of allosteric hubs are used to generate ligand-agnostic coupling values, whereas lists of ligand-exclusive allosteric hubs derived from molecular dynamics simulations are used to calculate dopamine or bromocriptine-specific coupling values. In each case, dynamic cross-correlation matrices (DCCMs) are generated for the list of hubs using the lowest 20 normal modes which has been shown^39,40^ to correspond to large-scale, concerted motions. The sum total of the dynamic cross-correlations between pairs of allosteric hubs serve as the coupling value.

